# Heterogeneity of transposon expression and activation of the repressive network in human fetal germ cells

**DOI:** 10.1101/364315

**Authors:** Boris Reznik, Steven A. Cincotta, Rebecca G. Jaszczak, Leslie J. Mateo, Joel Shen, Mei Cao, Laurence Baskin, Ping Ye, Wenfeng An, Diana J. Laird

## Abstract

Epigenetic resetting in germ cells during development leads to the de-repression of transposable elements (TEs). piRNAs protect fetal germ cells from potentially harmful TEs by targeted destruction of mRNA and deposition of repressive epigenetic marks. Here we provide the first evidence for an active piRNA pathway and TE repression in germ cells of human fetal testis. We identify pre-pachytene piRNAs with features of secondary amplification that map most abundantly to L1 family TEs. We find that L1-ORF1p expression is heterogeneous in fetal germ cells, peaks at mid-gestation and declines concomitantly with increasing levels of piRNAs and H3K9me3, as well as nuclear localization of HIWI2. Surprisingly, following this decline, the same cells with accumulation of L1-ORF1p display highest levels of HIWI2 and H3K9me3, whereas L1-ORF1p low cells are also low in HIWI2 and H3K9me3. Conversely, earlier in development, the germ cells lacking L1-ORF1p express high levels of the chaperone HSP90a. We propose that a subset of HSP90a-armed germ cells resists L1 expression, whereas only those vulnerable L1-expressing germ cells activate the PIWI-piRNA repression pathway which leads to epigenetic silencing of L1 via H3K9me3.

## Introduction

Tasked with the vital role of species propagation, germ cells must be able to faithfully transmit heritable unis of information between generations. A potential risk to genome integrity arises in the developing embryo as primordial germ cells (PGCs) undergo epigenetic reprogramming, characterized by DNA demethylation and global resetting of histone marks (Gkountela et al. 2015; Guo et al. 2015; Tang et al. 2015; 2016). Rearrangement of the epigenetic landscape primes germ cells for meiotic entry in fetal ovaries, and mitotic arrest and differentiation in fetal testis (Messerschmidt et al. 2014), yet risks reactivation of transposable elements (TEs) which are normally silenced. The greatest threat comes from retrotransposons, which comprise ~40% of the human genome (Kazazian and Moran 2017). Though most TEs are inactive and nonfunctional, some remain capable of retrotransposition, mostly stemming from the evolutionarily-young long interspersed element type 1 (L1), SVA, and Alu families (Kazazian and Moran 2017). L1-mediated integration of active TEs has been associated with over 100 disease mutations (Hancks and Kazazian 2016). Their activity is high, with new L1 and Alu integrations observed in ~ 1/100 and 1/20 human births, respectively (Hancks and Kazazian 2012), and even higher in mice, where new L1 insertions occur in 1/8 births (Richardson et al. 2017).

In germ cells, the activity of TEs is mollified by an evolutionarily conserved genome defense system made up of small RNAs and PIWI-family RNA-binding proteins collectively called the PIWI-piRNA pathway. Small (26–32 nt) RNAs, termed piRNAs (<u>P</u>IWI-<u>i</u>nteracting <u>RNA</u>s), complex with PIWI-family proteins to directly target retrotransposons for mRNA degradation in the cytoplasm, while also leading to nuclear silencing at endogenous genomic loci through DNA methylation and histone modifications (Weick and Miska 2014; Iwasaki et al. 2015). The PIWI-piRNA pathway is required for male fertility, and genetic mutants in this pathway invariably lead to meiotic catastrophe and failure to make mature sperm, while displaying increased expression of TEs along with decreased DNA methylation and loss of the repressive Histone 3 lysine 9 trimethyl (H3K9me3) mark at TE genomic loci (Aravin et al. 2007; Carmell et al. 2007; Aravin et al. 2009; Pezic et al. 2014). Elevated retrotransposition has also been observed in piRNA-deficient mice using a transgenic model for L1 mobilization (Newkirk et al. 2017). Germ cells deploy the PIWI-piRNA pathway at two stages during development: in prepachytene fetal germ cells and postnatally in pachytene stage germ cells (Czech and Hannon 2016; Iwasaki et al. 2015). In mice, the effector proteins PIWIL2 (MILI) and PIWIL4 (MIWI2) regulate transposons at multiple levels in fetal germ cells (Ernst et al. 2017). Nascent TE transcripts are processed into primary piRNAs, which are subjected to MILI-mediated amplification, generating antisense, secondary piRNAs. The latter are loaded onto MIWI2, resulting in nuclear localization and epigenetic silencing of TEs (Carmell et al. 2007). An array of co-factors is involved in the PIWI-piRNA pathway including the inducible chaperone Hsp90 (HSP90AA1/HSP90α in mammals), which assists in piRNA biogenesis, loading of piRNAs onto PIWI-family proteins, and transposon repression (Gangaraju et al. 2011; Ichiyanagi et al. 2014; Izumi et al. 2013; Specchia et al. 2010; Xiol et al. 2012).

Most studies of TE repression in developing mammalian germ cells have been conducted in mice. However it was recently shown that epigenetic reprogramming via global DNA demethylation and histone modification is conserved in human fetal cells (Gkountela et al. 2013; Guo et al. 2015; Tang et al. 2015; 2016), as is the accompanying activation of retrotransposon expression (Gkountela et al. 2015; Guo et al. 2015; Tang et al. 2015) and upregulation of PIWI-piRNA pathway genes (Tang et al. 2015). The expression of PIWI proteins and piRNAs has furnished evidence of an active piRNA pathway in adult human testis and in fetal ovaries (Ha et al. 2014; Roovers et al. 2015; Williams et al. 2015). Notably, piRNAs identified are derived from clusters and resemble the composition of mouse pachytene piRNAs rather than fetal pre-pachytene piRNAs which are largely transposon-derived (Aravin et al. 2007; 2008; Kuramochi-Miyagawa et al. 2008). In human fetal testis, piRNA expression has been difficult to detect due to low abundance (Williams et al. 2015; Gainetdinov et al. 2017), and reported localization of PIWIL2 (HILI) and PIWIL4 (HIWI2) in the cytoplasm does not support nuclear silencing by this pathway (Gomes Fernandes et al. 2018).

We provide evidence for an active PIWI-piRNA pathway in the human fetal testis. We establish timing of PIWI-family protein expression and observe a nuclear localization of HIWI2, suggesting a role in nuclear silencing. We identify TE-derived piRNAs with the signature of secondary biogenesis. L1 expression increases concomitantly with L1-derived piRNAs and PIWI-piRNA machinery in fetal germ cells during mid-gestation; conversely we find that L1 levels decline during development as levels of the repressive histone mark H3K9me3 increase. However, we observe surprising heterogeneity at a single cell level which suggests that only a subset of human fetal germ cells express L1 and deploy the PIWI-piRNA pathway to repress TEs via H3K9me3-mediated silencing, whereas other germ cells remain resistant.

## Results

### Dynamic expression and subcellular localization of PIWI homologs in male fetal germ cells

In mouse fetal testis, biogenesis as well as secondary amplification of pre-pachytene piRNAs depends upon MILI, which turns on at embryonic day (E) 12.5 (Aravin et al. 2008). In human fetal testis, we detected protein expression of HILI from gestational week (GW) 11 through 22, which corresponds to the period of sex-specific differentiation. HILI was found in the more differentiated population termed advanced germ cells (AGCs) marked by VASA (Figure 1A), as well as in coexisting PGCs defined by OCT4 (**Figure S1A**). Initially low levels of HILI at GW 11 increased through development, and protein was distributed throughout the cytoplasm as well as in puncta beginning at GW13, consistent with reported localization of MILI to pi-bodies, or intermitochondrial cement, in mice (Aravin et al. 2009). HIWI (PIWIL1) expression was not observed in fetal testes (**Figure S1B**), as expected based on its known adult-specific expression in human (Gomes Fernandes et al. 2018) and mouse (Deng and Lin 2002) (**Figure S1C**). HIWI2 was first detected in a small subset of AGCs at GW14, increasing from GW18 onward (Figure 1B). The subcellular localization of HIWI2 shifted from predominantly cytoplasmic before GW18 to nuclear in most AGCs by GW21-22 (Figure 1C). In mice, MIWI2 is mainly nuclear by E16.5-E17.5 (Aravin et al. 2008; 2009; Shoji et al. 2009; Kuramochi-Miyagawa et al. 2010), but remains cytoplasmic in fetal germ cells of piRNA pathway mutants including *Mili, Mael*, and *Mov10l1* (Shoji et al. 2009; Aravin et al. 2009; Zheng et al. 2010). Indeed, we observed a cytoplasmic to nuclear progression of MIWI2 from E16.5 to E17.5 in mouse fetal testes (**Figure S1D**). Nuclear localization of PIWIL4 homologs in other species requires piRNAs and cofactors such as HSP90a, and leads to transcriptional silencing of transposons at endogenous genomic loci (Ichiyanagi et al. 2014; Fu and Wang 2014; Iwasaki et al. 2015). Accordingly, the dynamic localization of HIWI2 between GW18 and 21 suggests that piRNAs are produced and transported to the nucleus for similar epigenetic functions in male fetal germ cells of humans.

**Figure 1:**
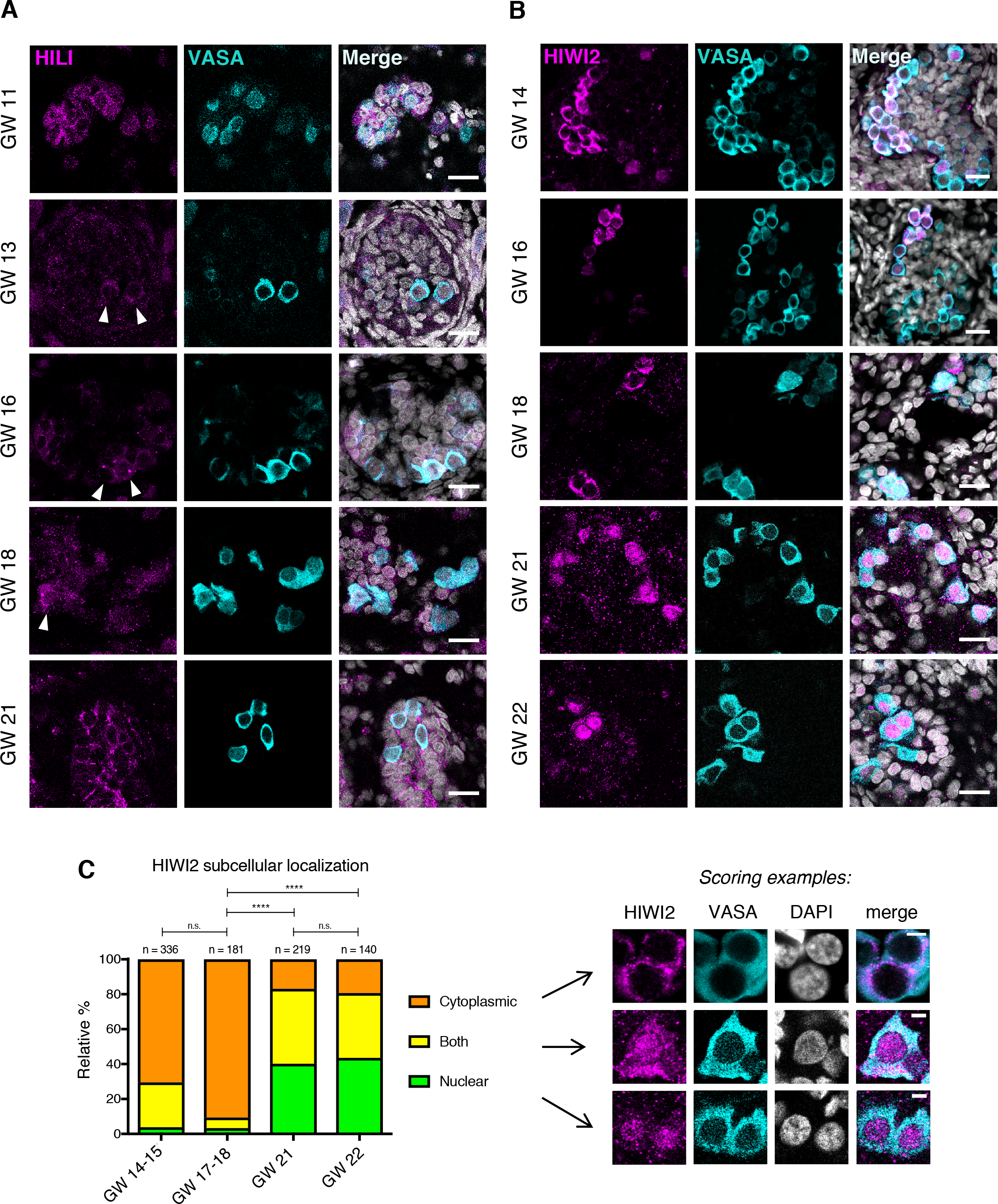
Expression of PIWI proteins in human fetal testis across development. (**A**) Expression of HILI and VASA at gestational week (GW) 11, 13, 16, 18, and 21, counterstained with DAPI (white, in merge images). Scale bars: 20 μm. Arrowheads indicate germ cells with HILI foci. A total of 12 embryos were analyzed across all time points. (**B**) Expression of HIWI2 and VASA at GW 14, 16, 18, 21, and 22, counterstained with DAPI. Scale bars: 20 μm. A total of 11 embryos were analyzed across all time points. (**C**) Relative frequencies of VASA^+^ germ cells with HIWI2 localization in cytoplasm, nucleus or both at indicated time points, with scoring examples at right. Two-tailed Fisher’s exact test was performed on “nuclear” and grouped “non-nuclear” categories across two time points; **** = p < 0.0001, n.s. = not significant. Scale bars: 5 μm.

### Evidence of transposon-derived primary and secondary piRNAs in fetal testis

We looked for evidence of piRNAs across timepoints spanning GW13 – 22. Small RNA sequencing was performed on bulk fetal testis tissue from eight samples at a depth of 35-40 million reads and processed according to our custom pipeline (Figure 2A). Trimmed 18-40 nt reads were first aligned to a curated list of human TE consensus sequences from the GIRI Repbase (Bao et al. 2015), and the remainder were aligned to the human genome (hg38). An overall alignment rate of >97% was obtained across all samples (**Table S1**). At all timepoints, the majority of reads (~75%) corresponded to miRNAs, with smaller fractions derived from tRNAs, snoRNAs, and protein coding transcripts (Figure 2B; **Figure S2A; Table S1**). Strikingly, less than 1% of reads mapped to Repbase, and these resolved into two size classes: 27 nt and 22 nt (Figure 2C). The 27 nt peak was highest at GW20, and likely corresponds to TE-derived piRNAs, which range from 26-32 nt in other species. Reads comprising the 22 nt peak showed extremely low sequence diversity, with the vast majority mapping to the L2C retrotransposon (data not shown). The number of Repbase-mapping reads was dynamic over development, with a maximum at GW20-21 (Figure 2D), corresponding to the window of HIWI2 nuclear translocation. Hallmarks of pre-pachytene piRNAs were identified in 26-32 nt Repbase-mapping reads, including a bias toward 1^st^ position uridine (Figure 2E, **S2B**) – a key signature of primary piRNA biogenesis (Iwasaki et al. 2015). Additionally, an adenine bias at the 10^th^ position at all timepoints with the exception of the earliest, GW13A (Figure 2E, **S2C**), suggests active secondary piRNA biogenesis, known as “ping-pong” amplification(Czech and Hannon 2016). Corroborating the specificity of Repbase-mapping reads, nucleotide distribution analysis on miRNA-annotated reads demonstrated a reduced, yet still apparent 5’ U bias (**Figure S2D**), similar to miRNAs in fetal ovary and adult testis (Williams et al. 2015). Another signature of ping-pong biogenesis, ten nucleotide overlaps between reverse complementary piRNAs, was detected in all samples to varying extents, with the exception of GW13A, and most strongly at GW20 (Figure 2F). Together these analyses identify small RNAs in human fetal testes with defining characteristics of pre-pachytene piRNAs, although at much lower abundance than found in mice or human fetal ovaries (Vasiliauskaitė et al. 2017).

**Figure 2:**
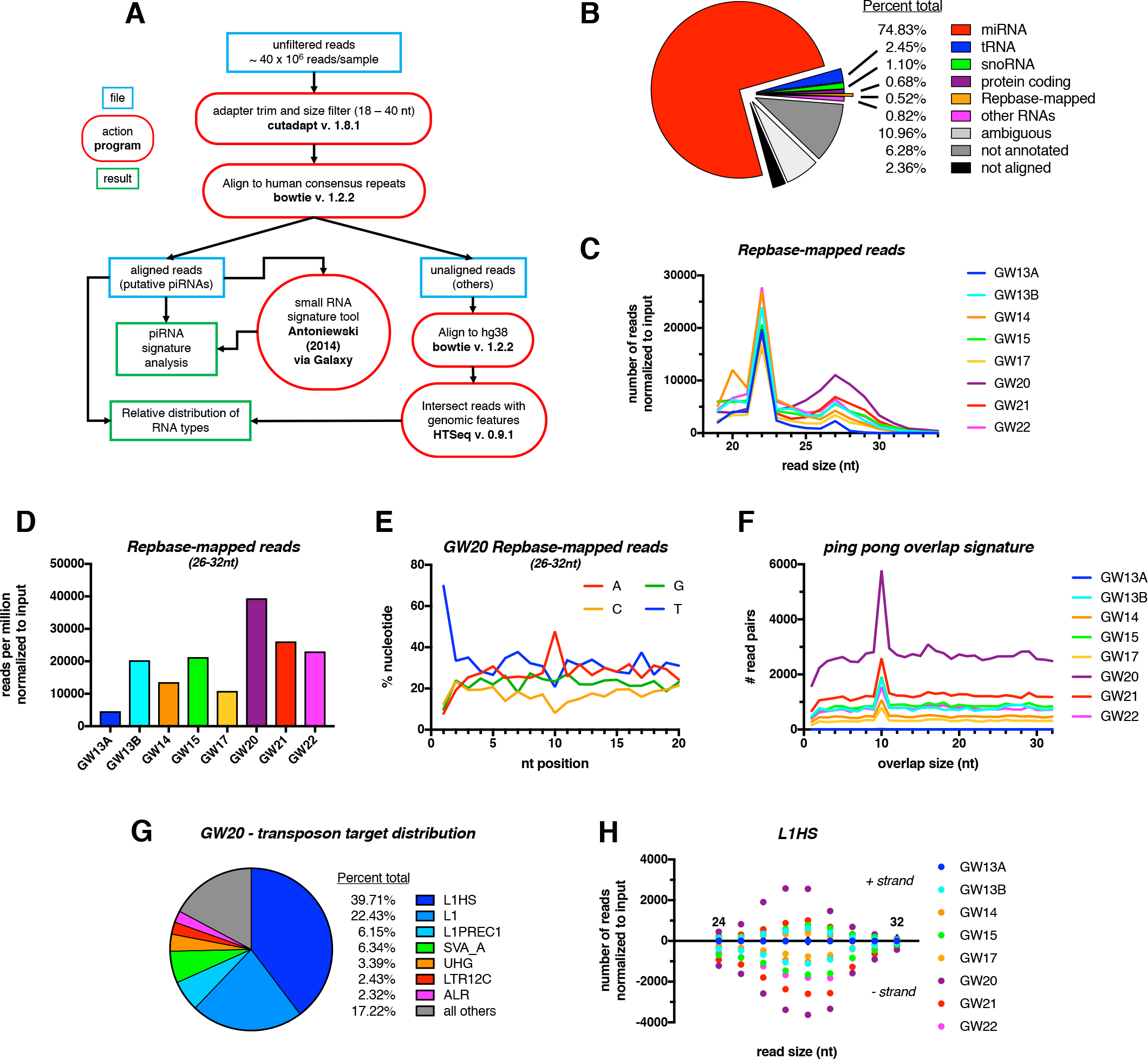
Identification of transposon derived piRNAs in the human fetal testis. (**A)** Schematic of small RNA-seq data analysis pipeline. (**B**) Relative percentages of different RNA subtypes present in gestational week (GW) 20 sample. (**C**) Size distribution of reads mapping to the Repbase database across all samples. (**D**) Normalized number of reads mapped to human transposon consensus sequences in Repbase database across all samples. (**E**) Relative nucleotide distribution at a given read position of pooled Repbase-mapping reads (26-32 nt) at GW20. (**F**) Ping-pong overlap signatures calculated on pooled Repbase-mapping reads (26-32 nt). (**G**) Top transposable elements mapped to by Repbase-mapping reads (26-32 nt) at GW20. (**H**) Distribution of read sizes mapping to the L1HS transposon across all samples. Positive and negative values represent reads mapping to the positive and negative strand, respectively.

Among Repbase-mapping reads, the L1 family was the most abundantly represented class of repeat elements at GW20 (Figure 2G). This is consistent with the derivation of piRNAs from actively-transcribed TEs, given that L1 elements are known to be active in humans (Hancks and Kazazian 2012). The evolutionarily youngest family (Smit et al. 1995), L1HS, was the most highly represented in all samples (with the exception of GW13A), followed by “L1” elements, which here represent the broad primate-specific L1PA clade (Kojima 2018) (**Table S2)**. Reads mapped to both forward and reverse strand of L1HS particularly in later stages (Figure 2H), consistent with progressive ping-pong amplification. Analysis of other highly-represented classes of TEs revealed different strand-specific biases, yet consistently peaked at GW20 (**Figure S2E-G**). We conclude that pre-pachytene piRNAs are produced and amplified in the human fetal testes, with a large fraction derived from TEs, similar to the developing mouse testis (Aravin et al. 2007; 2008).

### L1 transposons are expressed in AGCs coordinately with the transposon repression network at the single cell level

The prevalence of L1-derived piRNAs and possibility of piRNA-mediated nuclear silencing prompted examination of the relationship between transposons and the repression network in an unbiased, population-wide manner. We took advantage of a published single cell RNA-seq dataset from human fetal testis tissue (Li et al. 2017) to assess heterogeneity of transposon-derived and gene-coding transcripts. By mapping to repeat elements (based on RepeatMasker annotations) as well as the genome, we profiled expression of TEs as well as genes for each cell (Figure 3A). We focused on GW19 in light of the high level of piRNA expression and dynamic HIWI2 localization. t-distributed stochastic neighbor embedding (t-SNE) identified three distinct clusters corresponding to PGCs, AGCs, and somatic support cells based on known lineage- and stage-specific markers (Figure 3B; **S3A-D**). As anticipated, PGC and AGC clusters were distinguished by expression of *OCT4* and *VASA*, respectively (Figure 3C). The machinery of primary piRNA biogenesis (*MAEL* and *HILI*) was expressed in both PGCs and AGCs, though higher in the latter, whereas *HIWI2* mRNA was largely limited to AGCs (Figure 3C, D). This developmental delay in transcription of *HIWI2* as compared to *HILI* recapitulates the sequential expression of the proteins observed in fetal testis (Figure 1A, 1B). With few exceptions, TE expression was dramatically elevated in AGCs compared to PGCs, and absent in the soma (Figure 3C, D). The most highly expressed TEs in AGCs belong to the L1 clade, specifically L1HS, L1PA2, and L1PA3, corresponding with the abundance of L1-derived piRNAs (Figure 2G). Other highly expressed TEs include members of the Alu family, especially the young AluYa5 and AluYb8 subfamilies, known to be active in humans (Batzer and Deininger 2002). Paired correlation analyses between L1HS, and either *HILI* or *HIWI2* both yielded positive correlations (Figure 3E, F). Together this analysis shows that on a single cell level, the expression of the transposon repression network is upregulated in concert with the expression of transposons (Figure 3D).

**Figure 3:**
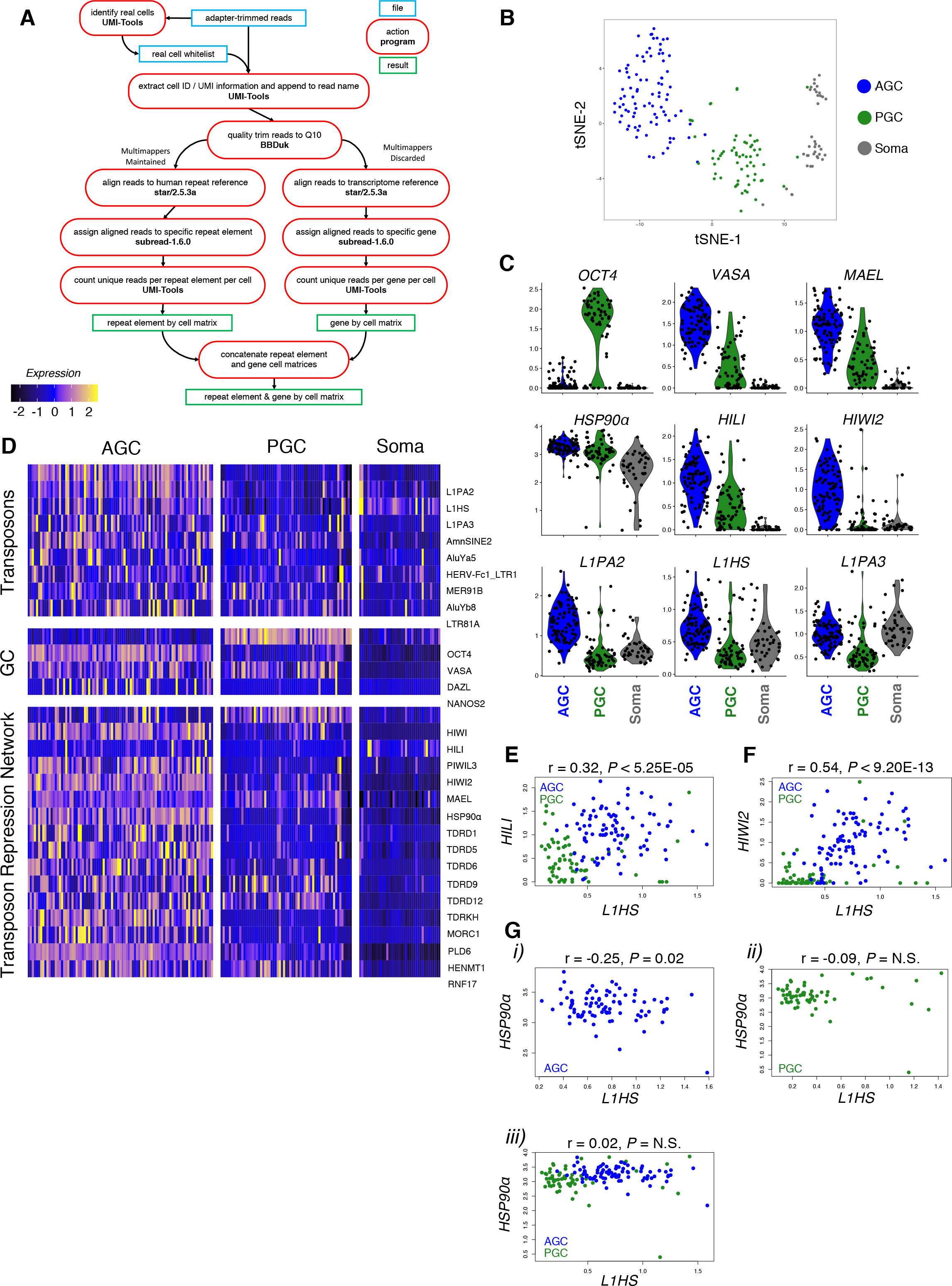
Single cell sequencing reveals dramatic upregulation of L1 family retrotransposons in advanced germ cells. (**A**) Single cell RNA-seq analysis pipeline to interrogate GW 19 male human germ cell sample for gene and transposon expression. (**B**) tSNE clustering on transcriptome reference was used to generate three distinct cell populations. (**C**) Violin plots showing expression of germ cell markers (top), transposon repression genes (mid), and L1 family retrotransposons (bottom) in AGCs, PGCs, and soma. (**D**) Heatmap displaying the most differentially expressed transposons sorted by myAUC score (top), and expression of germ cell markers (mid) and transposon repression genes (bottom). (**E-G**) Pairwise correlation analysis between L1HS and HILI (**E**) and HIWI2 (**F**) in GCs and HSP90α (**G**) in AGCs (i), PGCs (ii), and combined GCs (ii). Pearson’s correlation coefficient scores and p-values are listed above graph.

### Dynamic relationships between L1 and the repression pathway at transcript and protein level

We validated the correlation between L1 transcripts and the transposon repression network by immunostaining. Using a previously validated antibody (Rodić et al. 2014), we detected the L1-ORF1p RNA-binding protein in VASA^+^ AGCs starting at GW16, whereas earlier it was found sparsely and at low levels (**Figure S4**). L1-ORF1p was restricted to the cytoplasm and concentrated in granules, some of which co-localized with HILI (Figure 4A). However, L1-ORF1p granules were distinct from those containing HIWI2 (Figure 4B). As suggested by the heatmap in Figure 3D, a small subset of PGCs express high levels of L1 transcript but not the transposon repression network. We similarly observed rare instances of robust L1-ORF1p expression in OCT4^+^ PGCs (Figure 4C, **top panel**) and in cells lacking VASA and harboring low levels of OCT4 (Figure 4C, bottom panel). It is unclear whether these cells are transitioning between PGCs and AGCs or whether they will turn on the repression machinery, but this observation raises the question whether transposon protein expression precedes that of HIWI2 and piRNA biogenesis.

**Figure 4:**
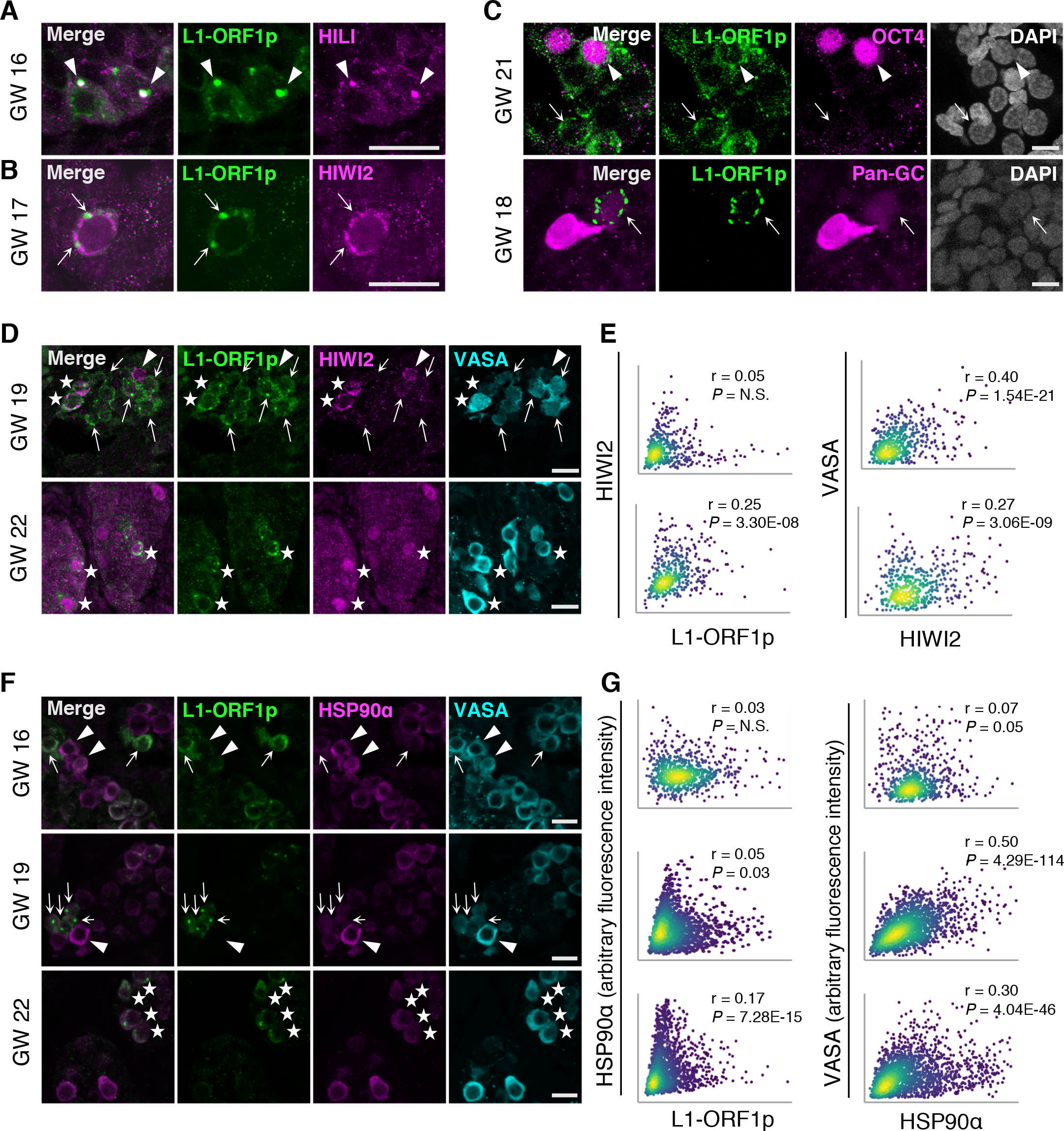
L1 expression in advanced germ cells. (**A**) L1 and HILI immunofluorescence in a GW16 human testis. Scale bars, 20 μm. Arrowheads indicate puncta of L1/HILI co-localization. (**B**) Expression of L1 and HIWI2 at GW17. Scale bars, 20 μm. Arrows indicate regions of mutually exclusive staining between L1 and HIWI2 puncta. (**C**) L1 expression at GW21 (top panel) and GW18 (bottom panel), co-stained with OCT4 or a pan-GC marker (OCT4 and VASA), respectively. (Top panel) Arrowhead indicates an L1^+^, OCT4^+^ germ cell, while arrows highlight L1^+^, OCT4-cell. (Bottom panel) Arrow indicates L1^+^ and pan-GC negative cell. Scale bars, 10 μm. (**D**) L1^+^ / HIWI2^low^ germ cells (arrows), HIWI2^high^ / L1^low^ germ cell (arrowhead), and double positive advanced germ cells (stars) in human fetal testis at GW19 and GW22. (**E**) XY scatter plots of HIWI2 vs L1 fluorescence intensities (left panel), and HIWI2 vs VASA (right panel), with Pearson’s correlation coefficients and corresponding p-values. For GW19, n = 526 VASA^+^ objects, counted from 3 stitched XY sections. For GW22, n = 472 VASA^+^ objects, from 2 stitched XY sections. (**F**) L1^high^ / HSP90α^low^ cells (arrows), HSP90α^high^ / L1^low^ cells (arrowheads), and double positive cells (stars) in human fetal testis at GW16-22. Scale bars 20 μm. (**G**) XY scatter plots of HSP90α vs L1 fluorescence intensities (left), and HSP90α vs VASA fluorescence intensities (right). Pearson’s correlation coefficients and corresponding p-values are noted. For GW16, n = 768 VASA^+^ objects, counted from 3 stitched XY sections. For GW19, n = 1809 VASA^+^ objects, counted from 4 stitched XY sections. For GW22, n = 2098 VASA^+^ objects, counted from 3 stitched XY sections.

Examination of the relative expression of L1-ORF1p, HIWI2, and VASA in AGCs revealed a more dynamic relationship than previously identified in the single cell RNA-seq analysis. The majority of L1-ORF1p expressing AGCs at GW22 co-expressed HIWI2 with significant correlation that approached the r value between HIWI2 and VASA (Figure 4D, E). Earlier, however, at GW19 we observed VASA^+^ cells with exclusive expression of either L1-ORF1p or HIWI2 and within the AGC population, correlation between the two proteins was lacking (Figure 4D, E). Compared to the strong correlation between L1 and *HIWI2* transcripts at GW19 (Figure 3F, **Figure S3E, F**), these results indicate a delay in the coordinated expression of HIWI2 and L1-ORF1p proteins during fetal development. It is likely that the VASA single-positive cells at GW22 represent newly differentiating PGCs that have not yet turned on HIWI2.

HSP90α is a chaperone with an inducible isoform enriched in germ cells, which has been implicated in transposon repression and piRNA biogenesis in flies and mice (Gangaraju et al. 2011; Ichiyanagi et al. 2014; Izumi et al. 2013; Specchia et al. 2010; Xiol et al. 2012). Levels of *HSP90α* transcript were high but variable in AGCs and PGCs (Figure 3C, D), with a negative correlation observed between *HSP90α* and L1HS in AGCs but not PGCs (Figure 3G). Consistent with this, immunofluorescence revealed that a significant number of AGCs at GW16 and GW19 exclusively expressed L1-ORF1p or HSP90α proteins but not both (Figure 4F, top and middle panels); accordingly, XY scatter plots for L1-ORF1p and HSP90α intensities displayed “arms” at the extreme end of each axis of AGCs with mutually exclusive expression (Figure 4G). This relationship changed by GW22, when most AGCs expressing L1-ORF1p also expressed HSP90α (Figure 4F, bottom panel), and a highly significant correlation was found (Figure 4G). Together these observations suggest that three AGC populations coexist at GW16—L1-high, HSP90α-high, and those low for both—and by GW22 a population of HSP90α single-positive AGCs persist, but the majority of L1^+^ cells also express moderate to high levels of HSP90α.

### Declining L1 expression and increasing H3K9me3 during fetal development

Thus far we have characterized expression of key proteins of the PIWI-piRNA pathway and confirmed expression of piRNAs and retrotransposons, yet it remains unclear if the PIWI-piRNA pathway represses retrotransposons in human fetal testis. Although we observed increasing nuclear localization of HIWI2 by GW21 (Figure 1C), examination of subsequent events is limited by availability of samples after GW22. We therefore turned to a xenograft model (Mitchell et al. 2010; Tharmalingam et al. 2018), in which human fetal testis implanted under the kidney capsule of immunocompromised mice will vascularize and continue to grow. Fragments of a single GW17 testis were grafted into separate mice and allowed to grow for 4 and 8 weeks (Figure 5A). While non-transplanted GW17 samples showed predominantly cytoplasmic localization of HIWI2 (~ 88% of cells), the frequency decreased in xenotransplanted samples after 4 or 8 weeks (41% and 60%; Figure 5B, 5C). This change in HIWI2 distribution suggests that grafted tissues develop to an equivalent of GW22 given the similar frequency of HIWI2 distribution, or else that HIWI2 does not egress from the cytoplasm in all germ cells. Quantification of L1-ORF1p levels demonstrated a significant, concomitant decrease over four weeks following transplant, with levels decreased by ~50% in individual AGCs (Figure 5D) and ~2/3 of VASA^+^ cells devoid of L1-ORF1p (Figure 5E). As most AGCs at this timepoint have entered mitotic arrest(Mitchell et al. 2010; Li et al. 2017) (data not shown), a decrease in the frequency of L1-ORF1p^+^ cells must arise through degradation of protein, in addition to locus-specific silencing or transcript degradation. To ask whether the decrease in L1-ORF1p levels correlates with an increase in DNA methylation over time, we performed immunostaining for 5-methylcytosine (5mC). While robust methylation was detected in surrounding somatic cells, we were unable to detect 5mC in VASA^+^ AGCs at GW17-22 or in xenografted tissues (Figure 5F, **S5**). This is consistent with previous reports of low global DNA methylation levels at GW26(Guo et al. 2017), suggesting that remethylation occurs at later timepoints.

**Figure 5:**
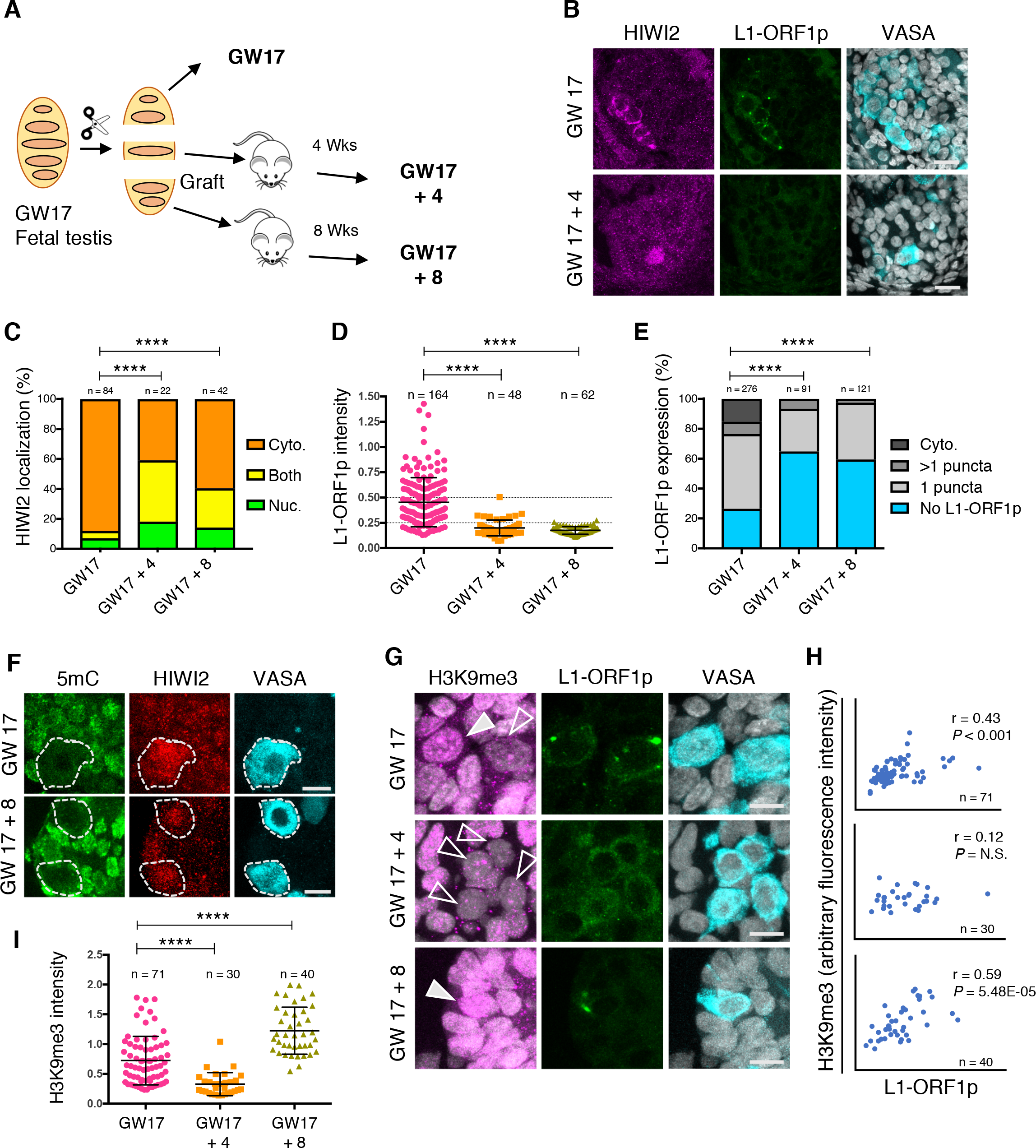
Evidence for PIWI-piRNA pathway activity in fetal testis. (**A**) Schematic of xenotransplant model. (**B**) Immunostaining for HIWI2, L1-ORF1p, and VASA (shown in merge with DAPI) at GW17 (no xenograft; top) and GW17 + 4 weeks xenograft (bottom). Scale bars: 20 μm. (**C**) Quantification of HIWI2 subcellular localization in VASA^+^ cells of xenografted testes, with scoring and statistics as in Figure 1C. (**D)** Quantification of L1-ORF1p intensity normalized to VASA; Mann-Whiney *U* tests were used to test significance. (**E**) Manual scoring of L1-ORF1p expression in VASA^+^ cells grouped into one of four categories: cytoplasmic staining, punctate cytoplasmic staining (either one puncta, or multiple puncta), or no L1-ORF1p expression. Two-tailed Fisher’s exact test was performed on the categories “no L1-ORF1p” vs all grouped “L1-ORF1p-expressing” cells across two time points to determine significance. (**F)** Immunostaining for 5mC, HIWI2, and VASA in fetal testis at GW17 and xenografts. Scale bars: 10 μm. (**G**) Immunostaining for H3K9me3, L1-ORF1p, and VASA (shown in merge with DAPI) at GW 17 and xenografts. Filled and empty arrowheads denote high and low H3K9me3, respectively. Scale bars: 10 μm. **(H)** XY scatter plots of H3K9me3 and L1-ORF1p fluorescence intensities with Pearson’s correlation coefficients, p-values, and number of cells counted. **(I)** Quantification of H3K9me3 intensity normalized to VASA, measured in Volocity. Mann-Whiney *U* tests were used to test significance. For all graphs, **** = p < 0.0001; n.s. = not significant; “n” represents number of individual cells counted per time point.

As piRNA-guided deposition of H3K9me3 marks has emerged as an additional mechanism for transcriptional silencing of retrotransposons (Pezic et al. 2014), we examined this repressive histone modification by immunofluorescence. Many VASA^+^ cells harbored few or no H3K9me3 puncta at GW17 and GW17+4, however we observed a significant increase in AGCs by GW17+8 (Figure 5G, I). Parallel trends of decreasing L1-ORF1p and rising H3K9me3 repressive mark would seem to be functionally related. However, unexpectedly, on a per cell basis, we observed that high levels of H3K9me3 were associated with high levels of L1-ORF1p (Figure 5H). Thus, we conclude that the developmental trends of L1-ORF1p and H3K9me3 in human AGCs parallel those described in mouse at the population level, but that our analysis of single cells opens up the possibilities that nuclear silencing operates independently of L1 protein degradation, or that some level of L1 protein persists in human AGCs. Our results also suggest that TE nuclear silencing via H3K9me3 is exclusively activated in the subset of AGCs that initially express high levels of L1 protein.

Our observations raise the possibility that L1 expression precedes that of the repressive pathway. Outstanding questions include the means by which the PIWI-piRNA network becomes activated as well as how TEs remain adequately repressed in PGCs given their demethylated state. The arginine methyltransferase PRMT5 has been implicated in TE repression in mouse PGCs by depositing repressive H2A/H4R3me2 marks on retrotransposons (Kim et al. 2014) and modification/activation of PIWI-family proteins (Vagin et al. 2009). Intriguingly, at GW13 when PGCs outnumber AGCs, PRMT5 could be detected in the cytoplasm and the nucleus, whereas it was exclusively cytoplasmic by GW17 (**Figure S6)**. In mouse male germ cells, such a nuclear to cytoplasmic translocation is observed by E11.5, signifying the transition from PRMT5’s role in histone modification to methylation of PIWI proteins (Kim et al. 2014; Vagin et al. 2009). Together, these results indicate that conserved mammalian mechanisms regulate TEs in the human testes. However, our analysis at the single cell level suggests that TE expression, and the resulting activation of the PIWI-piRNA pathway, occur in a subset of germ cells whereas others are resistant.

## Discussion

In preparation for gametogenesis, PGCs undergo epigenetic reprogramming which removes silencing marks that normally suppress TEs and leaves the germline vulnerable to DNA breaks or transposition. Our characterization of L1 expression in human fetal germ cell development (Figures 3, 4, 5, **S4**) underscores the need for the PIWI-piRNA pathway to protect germ cells from TE activity and maintain germline integrity. In this study we describe the dynamics of L1 expression in germ cells of the human fetal testis and provide evidence in support of an active TE repression network including key PIWI proteins, pre-pachytene piRNAs, and critical cofactors, such as HSP90α.

The fetal PIWI-piRNA pathway in mice is defined by PIWI family protein expression, TE-derived piRNAs, and piRNA-dependent deposition of repressive epigenetic marks (Ernst et al. 2017). We demonstrate conservation of these features in human fetal testis by characterizing: 1) composition, temporal expression pattern, and sub-cellular localization of PIWI-family proteins (Figure 1), and 2) the presence of transposon-derived piRNAs with a secondary amplification signature (Figure 2). In mouse germ cells, MIWI2 is expressed from E15 - P3 (Aravin et al. 2008), during which time it is required for deposition of the epigenetic silencing marks 5mC (Aravin et al. 2008; Kuramochi-Miyagawa et al. 2008) and H3K9me3 (Pezic et al. 2014) in order to fully deplete L1-ORF1p by P2 (Aravin et al. 2009). Similarly, we observed a coincident decline in the expression of L1-ORF1p and rise in the levels of H3K9me3 in AGCs from GW17 onward (Figure 5, **S4**). Given the limitations of working with human clinical samples, we could not test whether this was directly attributable to the PIWI-piRNA pathway. However, our cumulative results suggest that the fetal PIWI-piRNA pathway restricts TE expression in human fetal germ cells.

Our study provides the most comprehensive characterization of the PIWI-piRNA pathway in the human fetal testis to date. In addition to confirming and expanding upon the reported expression of HILI and HIWI2(Gomes Fernandes et al. 2018), we observed dynamic cytoplasmic to nucler localization of HIWI2 between GW18 – 21 (Figure 1). Recent studies in human fetal ovary and adult testis showed that piRNAs were derived from genomic piRNA clusters (Ha et al. 2014; Roovers et al. 2015; Williams et al. 2015). Previous efforts to identify piRNAs from fetal testis were confounded by low abundance, dearth of samples, and conflicting analyses (Williams et al. 2015; Gainetdinov et al. 2017). Here we characterized piRNAs from 8 fetal testes samples across a wide developmental range (GW13 - 22) and identified TE-derived piRNAs with a peak expression at GW20, which corresponds with the timing of nuclear HIWI2 translocation and further supports an active PIWI-piRNA pathway. This notion is corroborated by additional evidence including a correlation between high L1 transcript and lower expression of PIWI-piRNA pathway genes, including *HIWI2*, in boys at risk for infertility (Hadziselimovic et al. 2015). Additionally, mechanistic studies in human iPS cells showed that HILI can directly modulate L1 activity (Marchetto et al. 2013).

Human fetal germ cell development is marked by heterogeneity at the level of transcript, protein, and cell state. Whereas PGCs in mice acquire male differentiation markers and undergo mitotic arrest from E13.5 – E14.5 (Western et al. 2008), the transition from PGC to AGC in humans is spread across weeks hence both populations coexist (Gkountela et al. 2013). We observed heterogeneity between cells in the expression of transcripts for TEs and the repression network (Figure 3) and in levels of HIWI2 and HSP90α proteins (Figure 4). HSP90 is a highly expressed chaperone, constituting ~1% of cytoplasmic protein content (Buchner 1999). A germ cell specific, inducible isoform of HSP90 has been implicated in regulating the abundance of L1-ORF1p in mice (Ichiyanagi et al. 2014). Physical interaction between HSP90α and L1-ORF2p was recently revealed by proteomics (Taylor et al. 2018). In fetal testes at GW16-19, we observed reciprocal expression between HSP90α and L1-ORF1p, with AGCs positive for one or the other but not both proteins (Figure 4F); however, by GW 22, expression of L1-ORF1p was diminished overall (Figure 5C) and cells retaining L1-ORF1p co-expressed HSP90α (Figure 4F). A similar trajectory was found with HIWI2 and L1-ORF1p, in which AGCs at GW19 were primarily single positive, but expressed both proteins by GW22 (Figure 4E). Based on these observations, we propose a model whereby a subset of early AGCs marked by high levels of HSP90α at ĜW16 is resistant to the expression of transposons (Figure 6). Subsequently, AGC development diverges and HSP90α^low^ AGCs are susceptible to expression of L1-ORF1p, leading to upregulation of the PIWI-piRNA pathway and transposon repression through mRNA degradation and nuclear silencing via H3K9me3 (Figure 5G). Although we did not detect 5mC in AGCs in xenografted testes (Figure 5F), DNA remethylation at TE loci could be undetectable by immunofluorescence or else could occur later in development. By contrast, we propose that HSP90α^high^ AGCs are protected from initial transposon expression and may never express transposons or upregulate the PIWI-piRNA pathway. Our data suggest that suppression of TEs earlier in development in PGCs, prior to the expression of PIWI genes, could occur via other mechanisms such as PRMT5-mediated nuclear silencing (**Figure S6**), as previously shown in the mouse (Kim et al. 2014).

**Figure 6:**
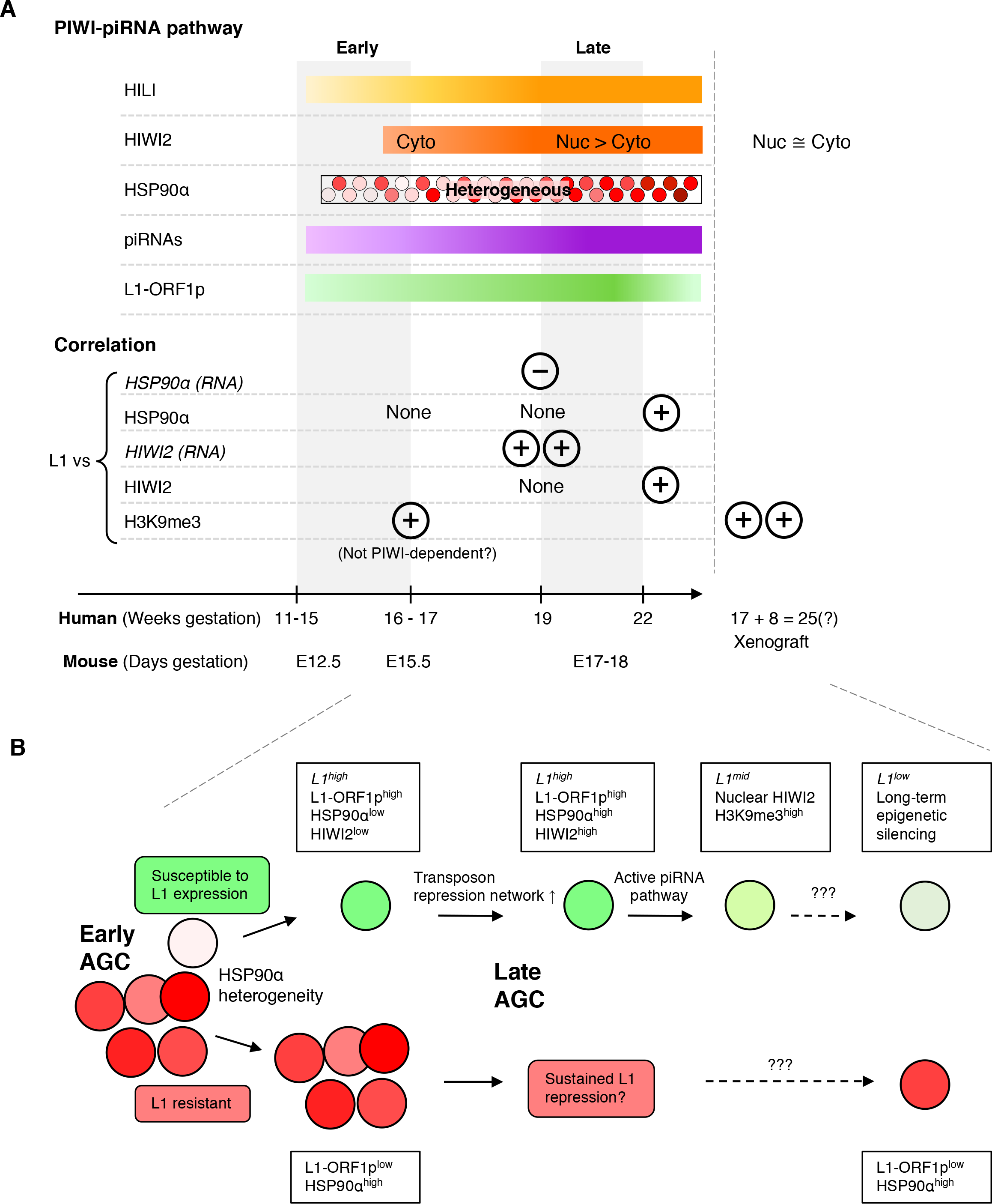
Coordinated transposon repression in GCs of the human fetal testis. **A**) (Top panel) Summary of the expression pattern of PIWI-piRNA pathway components and L1 protein in developing germ cells in the human fetal testis at indicated timepoints and in aged xenograft samples. (Bottom panel) Relationships between L1 expression (protein except where indicated) and expression of PIWI-piRNA pathway factors and H3K9me3 at indicated developmental stages. By GW22 many L1-expressing AGCs also express components of the repression network. (**B**) Model for transposon expression and repression in developing AGCs. See Discussion for details.

The PIWI-piRNA pathway is considered the immune system of the germline. In this paradigm, TEs are viewed as a threat to developing germ cells that require silencing to prevent their mobilization and subsequent genome disruption. Our single cell analyses reveal that the expression of TEs (Figures 3, 4) precedes that of the repression network. This premature exposure to TEs agrees with previous studies that have shown consistent L1 transcript expression from GW4 – 19 (Guo et al. 2015) where the earliest timepoint clearly predates the onset of the repression pathway. Could this early expression of TEs play a functional role during germ cell development? Recent studies argue in favor of this idea, by providing evidence that the L1 transcript is essential for early embryonic development in mice, by remodeling the epigenetic landscape (Jachowicz et al. 2017; Percharde et al. 2018). In addition, L1-ORF1p and L1-ORF2p were previously observed in a single GW18 testis (Ergün et al. 2004). As we detect the onset of HIWI2 and HSP90α later than the first L1-ORF1p (Figure 6), the initial expression of TEs may be required to “prime” the activation of the repression network. This question, and whether L1 plays any other functional roles during germ cell development will be the subject of future investigations.

## Acknowledgements

We are grateful to Brandon Sigamony, Daniel Nguyen, Suping Peng and other members of the Laird lab for their feedback and support as well as comments on the manuscript from Drs. Gerald Cunha and Miguel Ramalho-Santos.

## Author Contributions

DJL and BR designed the study; BR, LJM, RGJ, SAC, JS, and MC conducted experiments; BR, RGJ, SAC, PY, DJL and AW analyzed the data; DJL,BR, AW and SAC wrote the manuscript with edits and feedback from all authors.

## Competing Interests

None

## Funding

This work was funded by NIH R21ES023297, NIH R21ES023297 and R01ES028212 (to DJL), T32HD7263-31(to BR), 1F31HD096840 (to RGJ), K12DK083021 (to LB), 5T32HD007470 (to SAC), CIRM EDUC2-08391 (to LJM), and the Escher Fund for Autism (to DJL and WA).

## Methods

### Human sample collection and processing

Human fetal specimens were collected free of patient identifiers after elective termination of pregnancy (Committee on Human Research, University of California, San Francisco, IRB# 12-08813 and 16-11909). Age of specimen (referred to as GW – gestational week in text) was determined by heel-toe measurement. Gonads were left in RPMI 1640 media supplemented with L-Glutamine at 4°C from a few hours to overnight, and up to two nights for some samples. Specimens were either flash frozen for molecular biology analysis or fixed for immunofluorescent staining. For staining, tissues were fixed overnight in 2% PFA/PBS at 4°C with gentle agitation. Tissues were then washed, incubated in 30%, embedded and frozen in OCT solution (Tissue-Tek). Embedded tissues were sectioned at 8 μm thickness.

### Animals and xenograft model

WT embryos were generated by mating CD1 females with homozygous Oct4-ΔPE-GFP males (MGI:3057158, multiple-copy transgene insertion (Anderson et al. 1999). Embryos were dissected from timed matings and staged using anatomical landmarks from E16.5 – E18.5.

Immunodeficient CD1 nud/nud homozygous (Charles River) mice were used as host for xenograft experiments. Prior to surgery, human fetal testes were cut into four fragments and held in ice-cold RPMI 1640 medium, with one fragment set aside for fixation. Hosts were anesthetized and one fetal testis fragment was xenografted into the left kidney capsule of each host. Following surgery, animals were monitored daily for food and water intake and any obvious signs of stress. All mouse work was performed under the University of California, San Francisco, Institutional Animal Care and Use Committee guidelines in an approved facility of the Association for Assessment and Accreditation of Laboratory Animal Care International.

### Immunofluorescence

Slides were washed with PBS, permeabilized and then blocked for 1 hour at room temperature with blocking buffer (10% heat inactivated donkey serum in 1X PBS + 0.1% TX-100). Primary antibody (Table S3) incubation was performed overnight at 4°C, followed by washes and secondary antibody incubation (1:200) with DAPI (1:1000) for 1 hour at room temperature. Slides were washed and mounted with VECTASHIELD Mounting Medium (Vector Laboratories, Cat. No. H-1000). Some antibodies required antigen retrieval (see Table S3). Xenograft samples incubated with a mouse primary antibody required antigen retrieval and blocking with Mouse on Mouse blocking reagent (Vector Laboratories, Cat. No. MKB-2213).

### Immunofluorescence quantification

#### Automated fluorescence intensity quantification

Images were obtained and stitched from Keyence microscope (model BZ-X710) using a 20X objective. Background was subtracted on FIJI (Schindelin et al. 2012) using a rolling ball radius of 25 pixels and disabled smoothing. VASA^+^ objects corresponding to single cells were identified using Volocity version 6.3 using the following modules: “Find objects” in the VASA channel, “Close” objects x 2, “Fill holes in objects”, and separate touching objects at a size of 600 μm^2^. Next, VASA objects below and above size thresholds (< 599 μm^2^ and > 2405 μm^2^) were excluded. VASA^+^ objects were filtered for mean channel intensity > 45 and shape factor > 0.413. Objects were manually inspected and non-cellular objects were deleted. A variation of this protocol was used to find VASA^+^ objects from images acquired from the Leica Sp5 confocal microscope using a 63X oil objective.

#### Manual scoring for HIWI2 localization

Z-stack images were obtained on a Leica Sp5 confocal microscope using a 63X oil objective. Z-stacks were opened on FIJI and VASA^+^ cells were manually scored for HIWI2 subcellular localization. HIWI2 localization was compared to VASA and DAPI and assigned to either of three categories – nuclear (where the majority of HIWI2 appeared overlapped with DAPI staining), cytoplasmic (where HIWI2 staining was excluded from DAPI and overlapped with VASA) or both (where HIWI2 staining appeared equally distributed between the compartments). See Figure 1C for representative examples of cells scored for each localization phenotype.

### Small RNA-sequencing of human fetal testis

Samples were processed and sequenced by Quick Biology. Total RNA extractions were performed on fetal testis tissue using Trizol and quality control was performed using the Agilent Total RNA Nano Bioanalyzer kit. RNA integrity number (RIN) scores are provided in the table below. Libraries were prepared using the QIAseq miRNA Library Kit, and quality was assessed using the Agilent Bioanalyzer High Sensitivity DNA assay. Paired-end sequencing was performed on an Illumina HiSeq 4000.

| **Sample ID** | **Exact sample age** | **RNA QC (RIN score)** |
| --- | --- | --- |
| GW13A | 13 weeks | 6.3 |
| GW13B | 13 weeks | 6.3 |
| GW14 | 14 weeks | 6.9 |
| GW15 | 15 weeks | 6.6 |
| GW17 | 17 weeks | 5.9 |
| GW20 | 19 weeks, 4 days | 5.8 |
| GW21 | 20 weeks, 5 days | 5.2 |
| GW22 | 21 weeks, 5 days | 4.1 |

### Small RNA-sequencing data analysis

All samples were sequenced to a depth of approximately 35-40 million reads in order to meet the minimum read requirement for small RNA-seq data of 30 million as outlined by the ENCODE consortium (https://www.encodeproject.org/rna-seq/small-rnas/#restrictions). Raw reads in fastq files were trimmed to remove the adapter sequence (AACTGTAGGCACCATCAAT) using cutadapt version 1.8.1 (Martin 2011), and size filtered to 18 – 40 nt using cutadapt. FastQC (version 0.11.5) was utilized to broadly assess read quality. Trimmed reads were aligned to Repbase (version 22.11), a collection of consensus sequences of known human transposons curated by the Genetic Information Research Institute (Bao et al. 2015). Alignments were performed with bowtie version 1.2.2 (Langmead et al. 2009) using the following options: -v 1, -M 1, –best, –strata, allowing a maximum of one mismatch between the read and reference sequence, with multimappers being randomly assigned to only one location.

All reads that did not align to Repbase were subsequently aligned (with bowtie version 1.2.2 and same options) to the human genome (GRCh38). Aligned reads were intersected with genomic features and counted using the htseq-count function of HTSeq version 0.9.1 (Anders et al. 2015), with the following parameters: -f sam -t exon -s yes -i gene_type -m intersection-nonempty –nonunique all -a 0. The reference file containing genomic feature coordinates was a custom GTF file generated by merging the GENCODE 28 comprehensive gene annotation list with GENCODE 28 predicted tRNA genes. Feature counts for miRNAs, tRNAs, snoRNAs, and protein coding transcripts were combined with the number of Repbase-mapping reads to generate a breakdown of RNA subtypes (Table S1).

Repbase-mapped reads were further analyzed for piRNA signature, including a 5’ uridine bias and 10^th^ position adenine bias, by calculating the nucleotide frequency distribution per base position along the read for all pooled Repbase-mapped reads of a given sample. Read overlap signatures were calculated on the MISSISSIPPI Galaxy server using the “Small RNA Signatures” tool (Galaxy Version 3.1.0) (Antoniewski 2014). Transposable element annotations and transposable element-specific read size distributions were calculated directly from alignment files post Repbase mapping.

### Single-cell RNA-sequencing reanalysis

#### Bioinformatic Pipeline for single-cell RNA-seq data

Adapter trimmed reads were downloaded from Gene Expression Omnibus (GEO) (https://www.ncbi.nlm.nih.gov/geo/). FastQC (Conesa et al. 2016) was used for a broad quality control check. Using UMI-Tools (Smith et al. 2017) we determined cells from background noise using the cell ID / UMI information in read 2. Cell IDs / UMI information from read 2 was extracted and appended to the read name. Using BBDuk (Bushnell), reads were quality trimmed to Q10 (Mbandi et al. 2014; Mills 2014). For transcriptome alignment, we used default STAR/2.9.3a parameters (Dobin et al. 2013) to align to reference genome GRCh38.91 human transcriptome (Ensembl) with mitochondrial annotations added. To maintain the presence of multimappers in our human repeat reference alignment, the following STAR paramaters were used: –outFilterMultimapNmax 100, –winAnchorMultimapNmax 100, –outSAMmultNmax 100, –outFilterMismatchNmax 3 (Ge 2017). After specific gene or repeat element annotations were respectively assigned to the aligned files using Subread (Liao et al. 2014), UMI-Tools was used to count the unique reads per annotated gene/repeat element per cell. Count matrices of the human repeat reference alignment and transcriptome were concatenated, providing standard single cell RNA sequencing data on the transcriptome in conjunction with novel count data of repeat elements at a single cell level.

#### Clustering and differential gene expression analysis

Briefly, the concatenated count matrix was read into R/3.4.4 for analysis with the Seurat/2.1 suite of tools (Satija et al. 2015). Beginning with 196 cells from GW19, we filtered on a 0.06% mitochondrial gene expression threshold, leaving us with 191 cells. 24,432 genes and 1,225 repeat elements were expressed across all cells. Using Seurat, 782 variable genes were identified amongst the three defined clusters of somatic cells, primordial germ cells, and advanced germ cells. Transcriptome markers for the PGC and AGC populations were determined using the FindMarkers Seurat function and the “roc” test with a minimum cellular detection threshold of 0.25 for each population. Transposon expression data alone was appended as an assay to the Seurat object enabling us to ascertain which repeat elements were markers for each germ cell population.

## Supplemental Figure Legends

**Figure S1: Expression of PIWI proteins in human and mouse fetal testis.** (A) Immunofluorescence of GW18 human fetal testis to detect HILI and OCT4 expression. Merge is shown with DAPI co-stain. Arrows indicate HILI^+^/OCT^+^ germ cells. Scale bar: 50 μm. (B) Expression of HIWI in GW18 human fetal testis. Also shown is staining for AMH as a positive control (middle panel), and merged staining with DAPI (right panel). Scale bar: 100 μm. (C) MIWI staining of spermatogonia in adult mouse testis PND70. Similar staining and microscope acquisition settings were used as for the human sample in (B). Scale bar: 100 μm.(D) MIWI2 localization in murine fetal testis at E16.5 (top panel) and E17.5 (bottom panel) with subcellular localization of MIWI2 indicated. n = number of cells counted in four fetal testes across both timepoints. Arrow indicates a rare cytoplasmic MIWI2 germ cell at E17.5. Scale bars: 20 μm.

**Figure S2: small RNA-seq analysis of human fetal testis** (**A**) Relative percentages of different RNA subtypes present across all samples. (**B-C**) Relative nucleotide distribution at read position 1 (**B**) and read position 10 (**C**) for all samples, calculated on pooled Repbase-mapping reads (26-32 nt). (**D**) Control showing nucleotide distribution at read position 1 for miRNA-annotated reads only. (**E-G**) Distribution of Repbase-mapped read sizes mapping to the L1 (**E**), SVA-A (**F**), and L1PREC1 (**G**) elements across all samples, normalized to number of input reads. Positive and negative values represent reads mapping to the positive and negative strand, respectively.

**Figure S3: Validation of cluster identities in analysis of human fetal testis single cell RNA-seq** (**A-C**) Violin plots validating expression of additional germ cell markers (**A**), as well as somatic cell markers of both Sertoli cells (**B**) and Leydig cells (**C**). (**D**) tSNE plots depicting expression of germ cell markers and transposon repression genes in each cluster. (**E-F**) Pairwise correlation analysis between L1PA2 (**E**), and L1PA3 (**F**) against HILI and HIWI2 in the total population. Pearson’s correlation coefficient scores are listed above each graph.

**Figure S4: Developmental time course of L1 expression in human fetal testis** L1 and VASA immunofluorescence in GW11 - 22 human fetal testis, counterstained with DAPI (shown in merge). A total of 12 embryos were analyzed across all time points. Bottom panel shows no primary antibody control and background fluorescence in corresponding channels. Scale bars: 20 μm.

**Figure S5: No evidence for global DNA remethylation in AGCs**Immunostaining for 5mC, HIWI2, and VASA in fetal testis at indicated gestational ages **(A)** and in xenotransplant samples **(B)**. All scale bars indicate 20 μm except for bottom panel in **(A)** which is 50 μm.

**Figure S6: PRMT5 expression in human fetal testis** Immunostaining for PRMT5 in fetal testis at indicated gestational ages. Arrows indicate germ cells with nuclear accumulation while arrowheads indicate cells with cytoplasmic PRMT5. Scale bar: 50 μm.

